# PatchPerPixMatch for Automated 3d Search of Neuronal Morphologies in Light Microscopy

**DOI:** 10.1101/2021.07.23.453511

**Authors:** Lisa Mais, Peter Hirsch, Claire Managan, Kaiyu Wang, Konrad Rokicki, Robert R. Svirskas, Barry J. Dickson, Wyatt Korff, Gerald M. Rubin, Gudrun Ihrke, Geoffrey W. Meissner, Dagmar Kainmueller

## Abstract

Studies of individual neurons in the Drosophila nervous system are facilitated by transgenic lines that sparsely and repeatably label respective neurons of interest. Sparsity can be enhanced by means of intersectional approaches like the split-GAL4 system, which labels the positive intersection of the expression patterns of two (denser) GAL4 lines. To this end, two GAL4 lines have to be identified as labelling a neuron of interest. Current approaches to tackling this task include visual inspection, as well as automated search in 2d projection images, of single cell multi-color flip-out (MCFO) acquisitions of GAL4 expression patterns. There is to date no automated method available that performs full 3d search in MCFO imagery of GAL4 lines, nor one that leverages automated reconstructions of the labelled neuron morphologies. To close this gap, we propose PatchPerPixMatch, a fully automated approach for finding a given neuron morphology in MCFO acquisitions of Gen1 GAL4 lines. PatchPerPixMatch performs automated instance segmentation of MCFO acquisitions, and subsequently searches for a target neuron morphology by minimizing an objective that aims at covering the target with a set of well-fitting segmentation fragments. PatchPerPixMatch is computationally efficient albeit being full 3d, while also highly robust to inaccuracies in the automated neuron instance segmentation. We are releasing PatchPerPixMatch search results for ~30,000 neuron morphologies from the Drosophila *hemibrain* in ~20,000 MCFO acquisitions of ~3,500 Gen1 GAL4 lines.

**Code:** https://github.com/Kainmueller-Lab/PatchPerPixMatch

**Results:** https://pppm.janelia.org

## 1 Introduction

The recent release of a Drosophila melanogaster central brain connectome reconstructed from electron microscopy (*hemibrain* [21]), as well as a large resource of MCFO acquisitions of GAL4 line Drosophila central nervous systems (CNSs) [8,10] have paved the way for intersectional approaches [7] to sparsely and repeatably target a vast collection of individual neurons in the Drosophila CNS. To this end, a neuron of interest with known morphology, e.g. as reconstructed by electron microscopy, needs to be identified in MCFO acquisitions of two distinct GAL4 lines. Given such two GAL4 lines, a split-GAL4 line [13] can then be created, which expresses the positive intersection of the respective GAL4 expression patterns.

A key step in this approach is the identification of GAL4 line MCFO acquisitions that label a target neuron of interest. To this end, the NBLAST approach [6] has been shown to successfully identify neuron morphologies across imaging modalities [22]. However, NBLAST relies on curated reconstructions of target as well as source neuron morphologies as input. Such reconstructions have not been feasible to obtain at scale for GAL4 line MCFO acquisitions. An approach for neuron search that does not require reconstructions of individual neuron morphologies in MCFO images as input is Color-depth maximum intensity projection (MIP) search [11,1]. This approach operates on 2d projection images of MCFO channels, where it computes pixel-wise heuristic matching scores against projections of respective target neurons.

We describe here *PatchPerPixMatch*, an alternative fully automated approach that allows for efficient 3d search of neuron morphologies in GAL4 MCFO acquisitions. Our approach is based on PatchPerPix [9], a deep learning-based instance segmentation approach we have developed in previous work to tackle challenging properties specific to neurons in MCFO imagery, namely large spans of individual neuron instances, and overlaps of multiple instances as caused by partial volume effects. PatchPerPixMatch aims at piecing together a known target neuron morphology from PatchPerPix segmentation fragments. Thus, crucially, PatchPerPixMatch provides built-in robustness against false split errors in automated neuron instance segmentations.

We phrase the PatchPerPixMatch objective formally as a combinatorial optimization problem. We derive upper and lower bounds to the respective energy, and leverage these bounds to engineer an efficient solver. Thus we were able to run ~600 million PatchPerPixMatch searches, namely for ~30,000 target neuron morphologies reconstructed from the hemibrain [21] (version 1.2) in ~20,000 MCFO acquisitions of ~3,500 Gen1 GAL4 lines [10].

We are releasing the PatchPerPixMatch search results on *https://pppm.janelia.org*. Our code is available at *https://github.com/Kainmueller-Lab/PatchPerPixMatch*.

## 2 Method

Individual neurons in MCFO image data as released in [10] are infeasible to be reconstructed manually at scale. Respective suitable automated instance segmentation methods are scarce due to challenging characteristics specific to neurons in light microscopy, such as wide spans and complex shapes of neurons, as well as overlap of multiple neurons due to partial volume effects. In previous work, we have developed PatchPerPix [9], a deep-learning based instance segmentation method that is able to handle the above characteristics of neurons in light microscopy. PatchPerPix yields all neuron instances in an image in one go, yielding run-times that allow for processing MCFO data at scale.

PatchPerPix segmentations of neurons in MCFO image data are not to be considered correct reconstructions: Due to ambiguities in the data that often are hard to resolve locally even by eye, as well as the entailed scarcity of training data, false split- or false merge segmentation errors appear in many neurons. However, PatchPerPix segmentations do abundantly yield accurate reconstructions of large *fragments* of neurons.

The idea behind PatchPerPixMatch is to leverage PatchPerPix segmentations despite their inaccuracies, and thus allow for fully automated search of target neuron morphologies in MCFO data. To this end, PatchPerPixMatch searches for a target neuron morphology by determining a subset of PatchPerPix neuron segmentation fragments that jointly explain the target as well as possible, as illustrated in Figure 1. Thus PatchPerPixMatch achieves inherent robustness to false-split segmentation errors.

**Fig. 1:**
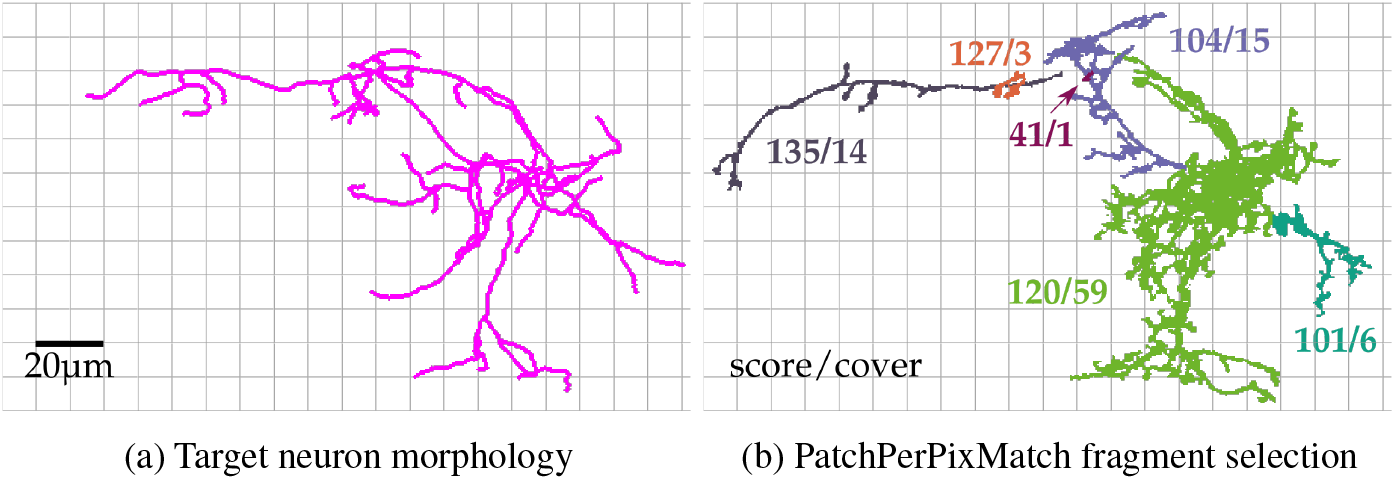
(a) Exemplary target neuron morphology (pC1e, see Resources Table in Figure 8), and (b) PatchPerPix instance segmentation fragments selected to cover the target neuron morphology (from an MCFO acquisition of the known matching GAL4 line R35C10). Each fragment yields a *score* that reflects how well it fits the target morphology locally, as well as a *coverage value* that reflects which percentage of the target the fragment may explain. Section 2.1 describes how we obtain scores and coverages per fragment, and how we derive a respective overall score for a selection of fragments. Sections 2.2 and 2.3 describe how we search for an optimal fragment selection for a given target neuron morphology.

The target neuron morphology is assumed to be given in skeletonized form, and in the same reference coordinate system as the segmentation fragments, which can e.g. be achieved via registration of the underlying imagery to a template brain [2]. In a first step, neuron segmentation fragments above a threshold size of 10*μm* (bounding box diagonal) are skeletonized [15], and NBLAST scores [6] are computed individually for each fragment vs. the target morphology. In a second step, PatchPerPixMatch aims at finding a subset of PatchPerPix fragments that jointly achieve *high NBLAST scores* and *cover large part of the target*, while also exhibiting *consistent colors* in terms of the three neuron-labelling channels of the underlying MCFO image. This second step is described in detail in Sections 2.1 to 2.3. Efficient processing of both hemispheres of the Drosophila brain is described in Section 2.4. While PatchPerPixMatch is inherently robust to false-split segmentation errors, it can, to some degree, also handle false-merge segmentation errors, as described in Section 2.5.

### 2.1 Objective

In formal terms, given a set of fragment IDs *F* ⊂ **N**, and a vector of respective indicator variables **x** = (*x*_1_, … , *x*_|*F*|_) ∈ {0, 1}^|*F*|^ that encodes fragment selection, the goal is to find a fragment selection that minimizes

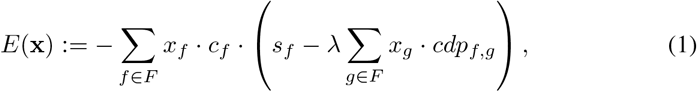

where *c_f_* measures which fraction of the target neuron morphology a fragment *f* covers, *s_f_* measures how well it scores in covering this fraction, and *cdp_f,g_* serves to penalize color differences between fragments (with weight *λ*). For convenient derivation of bounds to the objective (to follow in Section 2.2), the energy (1) can be re-written and refined as a sum over the points of a point cloud representation of the target, denoted via a set of point indices *T* ⊂ **N**:

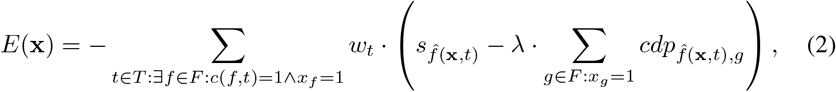

where *w_t_* weighs the influence of an individual target point, *c*(*f, t*) ∈ {0, 1} indicates if fragment *f* covers target point *t*, and 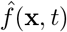 selects the best fragment to cover a target point *t* in case of multiple contenders.

In the following we give our specific definitions for each ingredient of (2), namely for weights *w_t_*, coverage *c*(*f, t*), best fragment 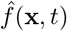, score *s_f_*, and color difference penalty *cdp_f,g_*.

#### Target Point Influence w_t_

Weights on target points are intended to increase the influence of stereotyped parts of a neuron’s morphology, i.e. parts with characteristic location as well as orientation, like e.g. wide-spanning projections, while downgrading the influence of e.g. small twigs in dendritic arbors, where orientation is less likely to be stereotyped [4]. To this end, given a skeleton representation of the target, we prune it by removing terminal branches that are shorter than a threshold length of 10*μm*, repeatedly for 3 iterations. Then, we assign a “branch length” to each remaining point of the target morphology, as follows: All points in terminal branches are assigned their respective branch’s length. Subsequently, the points that have already been assigned a length are ignored, and the procedure is iterated with the remaining morphology, until all points are processed. Finally, the point weights are normalized, such that 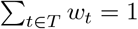.

#### Coverage c(f, t)

For an individual fragment *f*, we define the set of target points it covers, {*t ∈ T* : *c*(*f, t*) = 1}, as specified in Algorithm 1. The algorithm assumes a point cloud representation of the fragment as well as the target as inputs. Note that target- and fragment point cloud representations are expected to exhibit comparable sample point distances. We refer to the target point cloud as *P* := *p_t_* ∈ **R**^3^ : *t ∈ T*}, and to the point cloud for fragment *f* as *Q* := *q_i_* **R**^3^ : *i* ∈ {1, …, *N*}}.

Algorithm 1 considers the set *Q* of all fragment points jointly, with the desired effect of putting one large fragment at an advantage as opposed to many small fragments, as explained by the example sketched in Figure 2. Algorithm 1 restricts the number of target points a fragment point can cover to some maximum number to avoid over-coverage, as exemplified in Figure 3a. Over-coverage would otherwise be caused, e.g., by fragment end points that are closest to many target points within threshold distance. We empirically determined a restriction to six target points to yield visually plausible coverage. The *repeat* loop in Algorithm 1 avoids putting large fragments at a disadvantage as opposed to many small fragments, as exemplified in Figure 3b: E.g., a spurious side branch of a fragment could be closest to many target points, only a restricted number of which will be determined as covered. Our proposed looping gives the thus-excluded target points a chance to still be covered by other fragment points.

**Algorithm 1:**
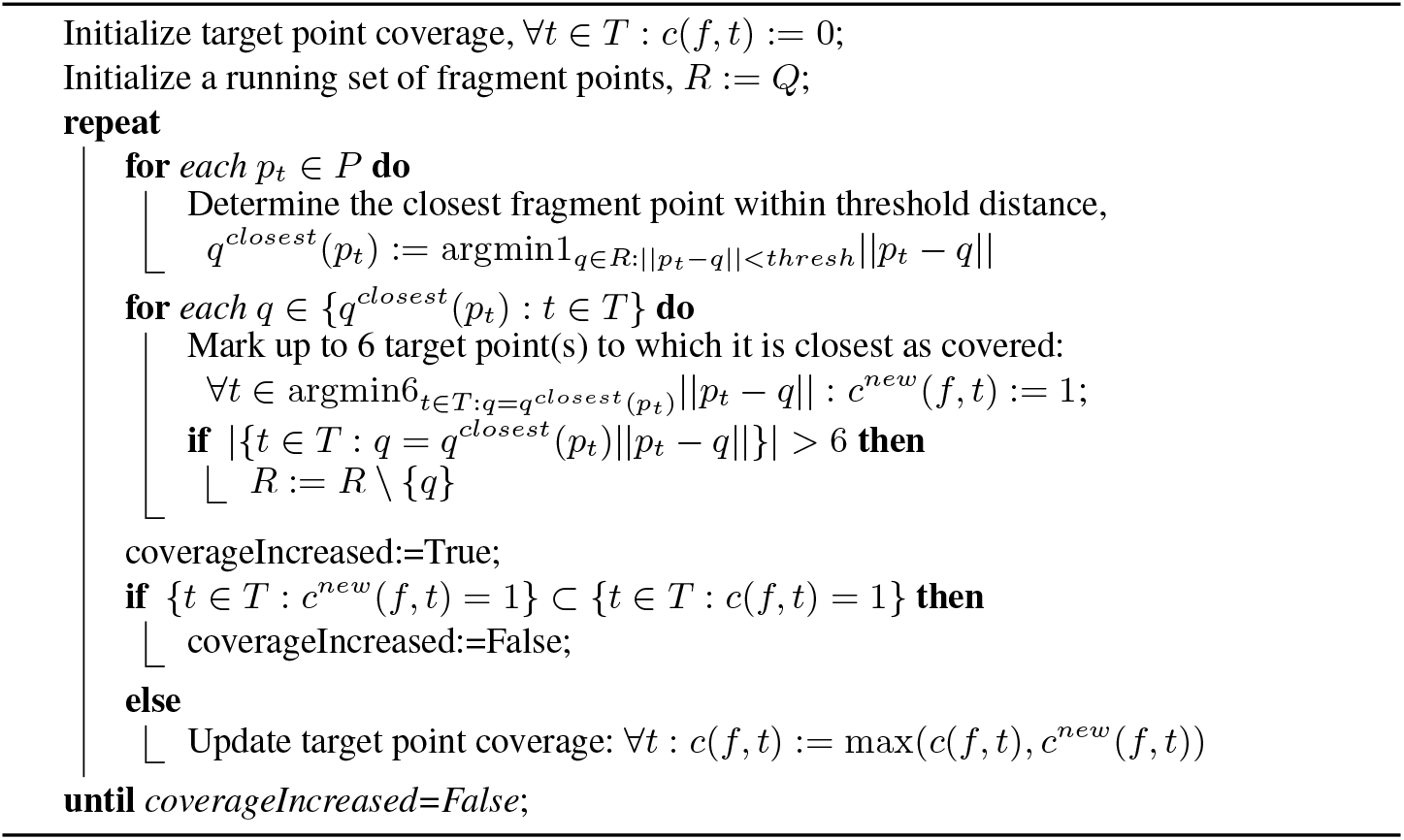
Target Point Coverage by a Fragment

**Fig. 2:**
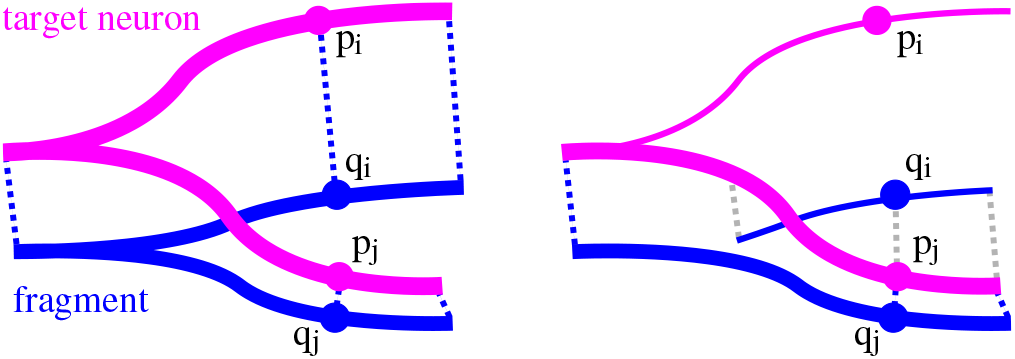
Algorithm 1, which serves for determining which portion of a target neuron morphology (pink) a neuron segmentation fragment (blue) covers, favors larger fragments: If a fragment is whole (left), *q_j_* covers *p_j_*, and *q_i_* covers *p_i_* because *p_j_* is already ”taken”. If instead the fragment is split into two parts (right), both *q_i_* and *q_j_* cover *p_j_* as it is the closest target point for both of them, and therefore the upper part of the target neuron will not be covered, indicated by thin (as opposed to bold) magenta line. (Blue dotted lines indicate desired matches, grey dotted lines indicate obsolete matches.)

**Fig. 3:**
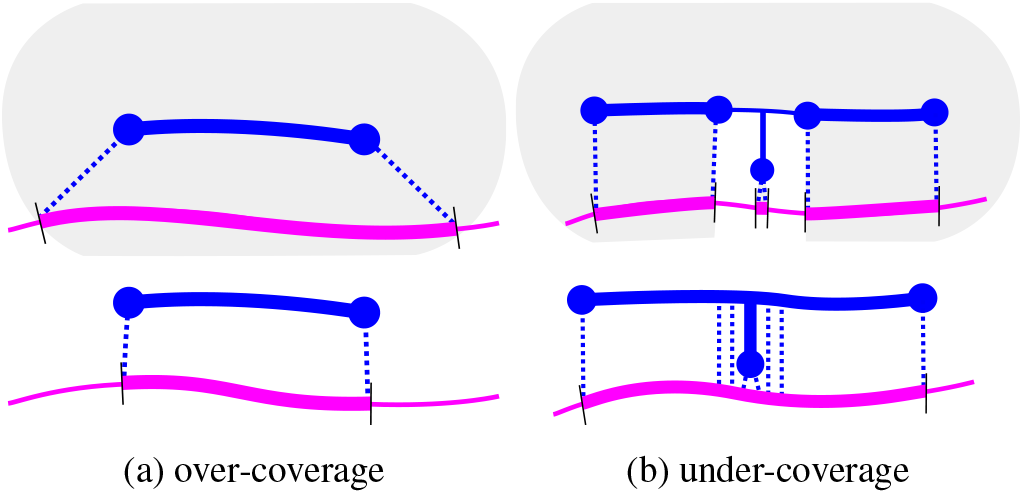
(a) Top: Without the constraints imposed by Algorithm 1, a fragment (blue) would cover an oversized portion of a target neuron, namely all target points within threshold distance (shaded area / bold magenta line). Bottom: Restricting the number of target points a single fragment point can cover results in coverage of an appropriately-sized target region. (b) Top: In case of a small side-branch, avoiding over-coverage as in (a) can lead to under-coverage, as many target points may have the same branch end-point as their closest point. Bottom: To avoid such under-coverage, Algorithm 1 loops over the point matching process, each time removing fragment points whose coverage had to be restricted in the previous iteration, until coverage does not increase any more.

#### Best Fragment 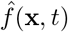

Given two fragments, the sets of target points they cover are not necessarily disjoint. To ensure that each target point contributes at most once to the objective (2), for each target point, we select one fragment that covers it ”best”. While seemingly straightforward to select the best-covering fragment for a target point by means of best NBLAST score, this choice has the drawback of favoring small fragments, as high NBLAST scores are abundantly obtained “by chance” by small fragments. Thus, to avoid false-positive solutions assembled from large amounts of small fragments, we instead employ as selection criterion the (weighted) amount of the target a fragment can explain, thus favoring larger, more characteristic fragments:

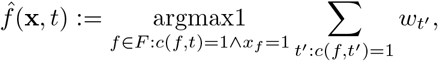

where we denote argmax1 := min(argmax(.)) for convenience to include the case of non-uniqueness of argmax.

#### Fragment Score s_f_

As an additional means to reduce the impact of small fragments (and finally only use them as “hole fillers”), we do not use the plain NBLAST score *nblast_f_* ∈ [-1, 1] of a fragment as score *s_f_* in (2). Instead, first, we cap the normalized NBLAST score at 0.5 to dampen the impact of small non-distinctive fragments that may easily receive very high scores. Second, we adjust the capped score according to the amount of target a fragment can (at most) explain, thus defining a positive, “coverage-adjusted” score, as follows:

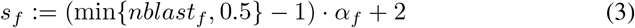

with

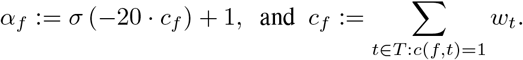

With this, *α_f_* approaches 1 for large fragments, thus yielding a capped and positive-shifted NBLAST score, min {*nblast_f_*, 0.5} * 1. Instead, for small fragments, *α_f_* approaches 1.5, thus yielding a score that is smaller than the capped and positive-shifted NBLAST score (as *α_f_* is multiplied to the *negative-shifted* NBLAST score).

#### Color Difference Penalty cdp_f,g_

A color is assigned to an individual fragment via K-Means clustering of the respective MCFO image voxels’ three channel values. We empirically set *k* = 2, and set the fragment color *col_f_* ∈ **R**^3^ to the mean of the larger cluster. Fragment colors for an individual MCFO acquisition are then normalized. We aim at a penalty that is small for differences in intensity, and large otherwise. More specifically, we follow the empirical observation that channel intensities within one neuron may fluctuate individually, though the channels that do fluctuate tend to do so in concordant direction. Hence we define the color difference penalty, based on color difference *cd_f,g_* := *col_g_* − *col_f_* ∈ **R**^3^ and a color difference threshold *cdt*, as follows:

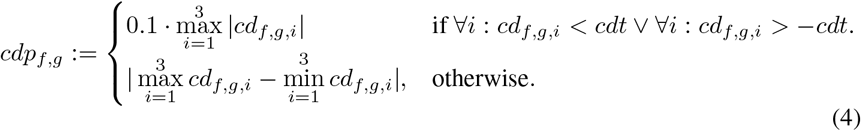

Thus, a small penalty is assigned if color channels fluctuate in concordant direction (where concordance is determined with slack *cdt*), whereas a larger penalty is assigned otherwise. The weight *λ* in (2) is set to adjust the color difference penalty for the number of selected fragments, and to weigh color difference penalties against scores. We empirically set 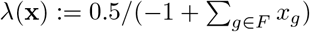.

Note that the PatchPerPixMatch energy (2), given our definitions of all of its components, can be modelled as a Markov random field (MRF), where nodes represent fragments, and binary labels represent fragment selection. However, the factorization of the respective probability distribution contains not just 2nd order terms (due to the color difference penalites), but also higher (> 2nd) order terms due to the dependency of best fragments 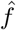 and weight *λ* on the overall fragment selection **x**. Consequently, generic efficient approximate solvers for 2nd order binary MRFs, like e.g. QPBO [14], do not apply.

### 2.2 Bounds

For efficient optimization of the PatchPerPixMatch objective (2), we define computationally cheap upper and lower bounds, as follows:

*Upper bound to the minimum energy:*

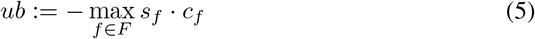

Our upper bound constitutes the minimal energy achievable with a single fragment solution.

*Lower bound to the minimum energy:*

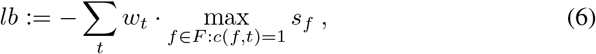

Our lower bound is obtained by picking the highest-scoring of all fragments per target point and ignoring the color difference penalty. This is a valid lower bound because

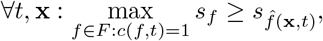

and furthermore, the color difference penalty is always positive, i.e., ∀*f, g* : *cdp_f,g_* ≥ 0.

*Lower bound to the energy of a candidate solution* **x**:

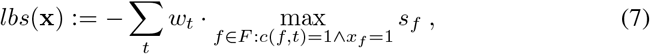

analogous to the lower bound to the minimum energy.

### 2.3 Solver

For efficient optimization of the PatchPerPixMatch objective (2), we first determine a set of candidate solutions heuristically via agglomerative clustering [20] of fragments by color difference. For each candidate solution, we then discard fragments whose total color difference penalty outweighs their score. We continue to the next candidate solution early if a lower bound (7) to the energy of the candidate surpasses a running upper bound to the minimum energy. The solver is specified in detail in Algorithm 2.

### 2.4 Efficient processing of both brain hemispheres

Due to fly brain symmetry, most target neurons may be found in either hemisphere of a segmented brain [12]. PatchPerPixMatch aims at finding the best match across hemispheres. To this end, we mirror the target neuron in the reference coordinate frame. For efficient processing of both hemispheres, we first determine the computationally cheap upper and lower bounds (5) and (6) for each hemisphere individually. If the lower bound determined for side I exceeds the upper bound determined for side II, side I can be discarded. Otherwise, we proceed with side I, and use the energy of its best solution as initial upper bound for side II.

**Algorithm 2:**
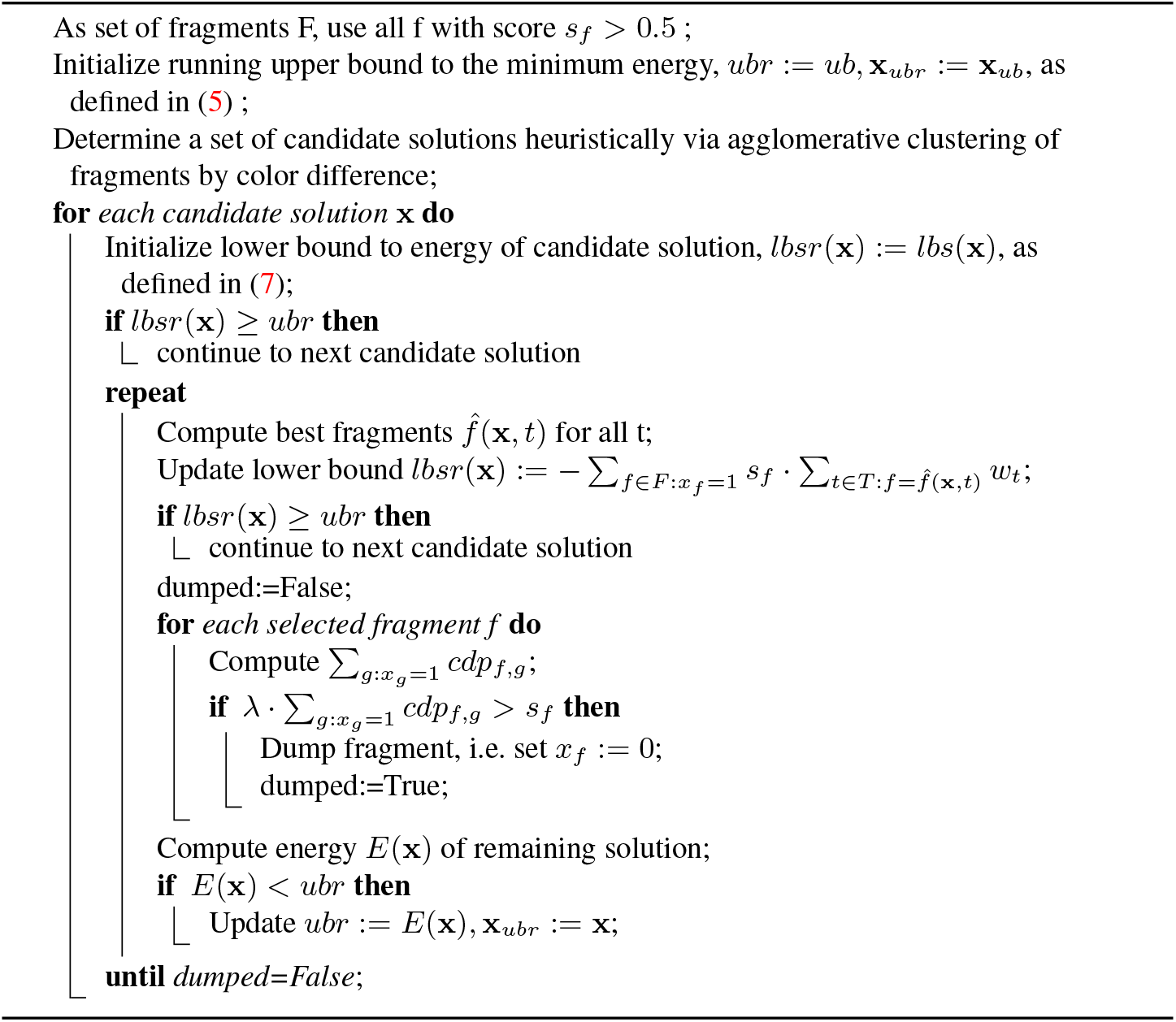
PatchPerPixMatch Solver

### 2.5 Robustness to false merge segmentation errors

PatchPerPixMatch naturally handles false split errors in the underlying segmentation by seeking a combination of segmentation fragments to best cover a target. False merge errors, on the other hand, may lead to false misses of matching neurons. This is mainly due to the fact that false merge fragments necessarily yield poor NBLAST scores, as a false merge fragment by definition cannot be a subset of any target morphology.

To nevertheless increase robustness of PatchPerPixMatch against false merge segmentation errors, we proceed as follows: We compute a second set of NBLAST scores, for which we prune each fragment to the part that lies within 20*μm* of the target. False merge fragments may thus yield decent NBLAST scores. We perform the whole PatchPerPixMatch pipeline again for this second set of NBLAST scores, where we re-use previously computed coverage values *c_f_* to avoid redundant computation.

Pruned fragments necessarily yield higher or equal NBLAST scores as compared to the original fragments. Hence PatchPerPixMatch energies of solutions obtained in the ”pruned” pipeline run cannot be directly compared with energies of solutions obtained in the ”vanilla” pipeline run. Thus we rank the lists of matches yielded by the two pipeline runs independently, in terms of PatchPerPixMatch energy. To yield one final ranked list while accounting for the fact that energies are not directly comparable across runs, we merge the lists from the two pipeline runs *by rank* (and not by energy). Specifically, we add 10.5 to the ranks yielded by the pruned run to achieve uniqueness and to slightly favor the vanilla run, then sort the merger of both lists by rank, and finally remove duplicate matches with lower ranks from the merged list.

## 3 Results

We ran PatchPerPixMatch for ~30,000 target neuron morphologies from the hemi-brain [21] version 1.2, selected by means of size and quality tags via the *neuPrint* tool [5] publicly available at *https://neuprint.janelia.org*. We searched for these target neuron morphologies in ~20,000 MCFO brain acquisitions of sparse and medium density Generation 1 GAL4 lines (Gen1 MCFO Phase 1 Categories 2 and 3 as published with [10], available at *https://gen1mcfo.janelia.org*, registered to the JRC2018 Unisex template [2]).

### Bulk quantitative analysis

We evaluated the accuracy of PatchPerPixMatch quantitatively on 10 target neuron morphologies and respective 47 known matching GAL4 lines [17,19,18], as listed in Appendix A, Figure 8. For each known matching GAL4 line, we determined two different ranks: (1) The rank of the best-matching MCFO acquisition of the GAL4 line among all ~20,000 searched MCFO acquisitions (termed *sample rank*), and (2) the rank of the best-matching MCFO acquisition of the GAL4 line among all ~3,500 best-matching MCFO acquisitions per GAL4 line (termed *line rank*).

The median sample rank of an MCFO acquisition of a known matching line is 23. The median respective line rank is 18. I.e., for half of the known matching lines, the best-matching respective MCFO acquisition ranks at or above 23 among all ~20,000 samples, and at or above 18 among all ~3,500 best-matching samples per line. In terms of sample ranks, best-matching MCFO acquisitions of the known matching lines are ranked among the top 1000, top 500, top 150, and top 50 samples for 42, 40, 36, and 26 of the 47 known matches, respectively. In terms of line ranks, best-matching MCFO acquisitions of the known matching lines are ranked among the top 500, top 150, and top 50 lines for 42, 38, and 30 of the 47 known matches, respectively.

### Quantitative analysis for the neurons pC1d and pC1e

For the neuron pC1d, a set of known matching GAL4 lines has been determined via exhaustive behavioral screen of nearly all Janelia (”R”) lines [16] and thus exhibits an exceptional level of completeness. Furthermore, an overlapping, yet not identical set of matching lines is also known for the morphologically similar neuron pC1e. These circumstances exclusively allow for a quantitative assessment of false PatchPerPixMatch hits. Note that the ”VT” lines were not included in the behavioral screen and were hence excluded from our respective analysis. Figure 4 shows exemplary morphologies of the pC1d and pC1e neurons. Note the distinctive arbor present in pC1d but not in pC1e.

**Fig. 4:**
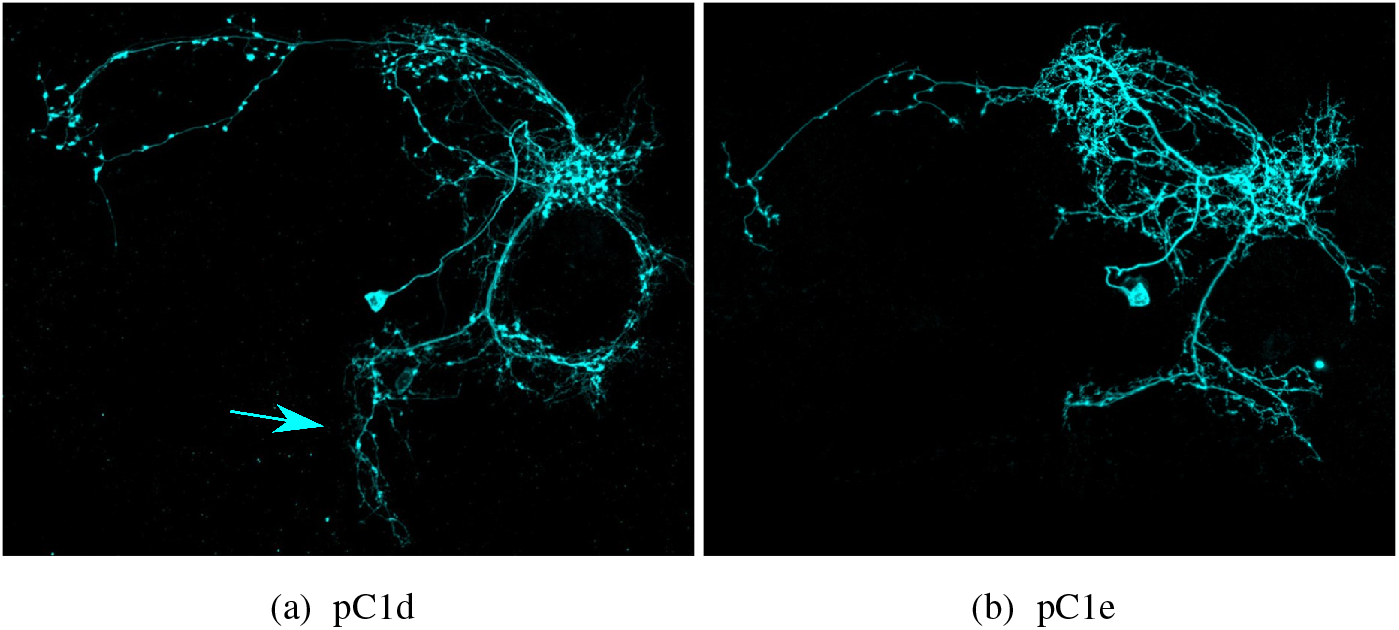
Exemplary pC1d and pC1e neurons (excerpts from Figure 4 – figure supplements 3D and 4D of [17]). pC1d features a distinctive arbor, see arrow.

Appendix B, Figure 9 lists the PatchPerPixMatch line- and sample ranks of the assessed known matching lines for pC1d and pC1e. Appendix B, Figure 10 lists the first 50 PatchPerPixMatch hits for pC1d as well as for pC1e. All non-confirmed Janelia line hits were inspected visually and deemed as either plausible, not convincing, or too faint or crowded to judge. For pC1d, out of the 27 Janelia line samples among the first 50 PatchPerPixMatch hits, 16 are samples from confirmed matching lines, 4 are confirmed matches for the morphologically similar pC1e, 2 are suspected matches of the morphologically similar pC1e, 2 are too crowded or faint to judge, and 3 do not look plausible. For pC1e, out of the 28 Janelia line samples among the first 50 PatchPerPixMatch hits, 17 are samples from confirmed matching lines, 2 look plausible albeit not confirmed, 3 are too crowded or faint to judge, and 6 do not look plausible. Two lines that are known to contain pC1e but not pC1d, namely R60G04 and R60G08, are ranked at line ranks 1 and 3 for pC1e. Respective false hits for pC1d rank comparatively lower, namely at line ranks 10 and 12, with 4 and 5 known matching lines, i.e. true hits, ahead of them, respectively.

### Qualitative analysis

We inspected PatchPerPixMatch hits visually to assess its potential to reveal matches to end users, as well as to determine typical causes for falsely missing known matches. Figure 5a showcases a true positive PatchPerPixMatch hit for a relatively sparse MCFO sample, which is easily visible by eye in a maximum intensity projection (MIP) of the MCFO image (see row 1). Figures 5b and 5c showcase true positive hits for more challenging MCFO acquisitions, namely for densely clustered and dim neuronal morphologies. These are hardly visible by eye either in the MIP of the MCFO image (row 1), or in color-depth MIPs [11] of the MCFO channel that best labels the matching morphology (rows 4 and 5). However, the hits are revealed by masking the MCFO image with the instance segmentation fragment selection determined by PatchPerPixMatch (rows 2 and 3).

**Fig. 5:**
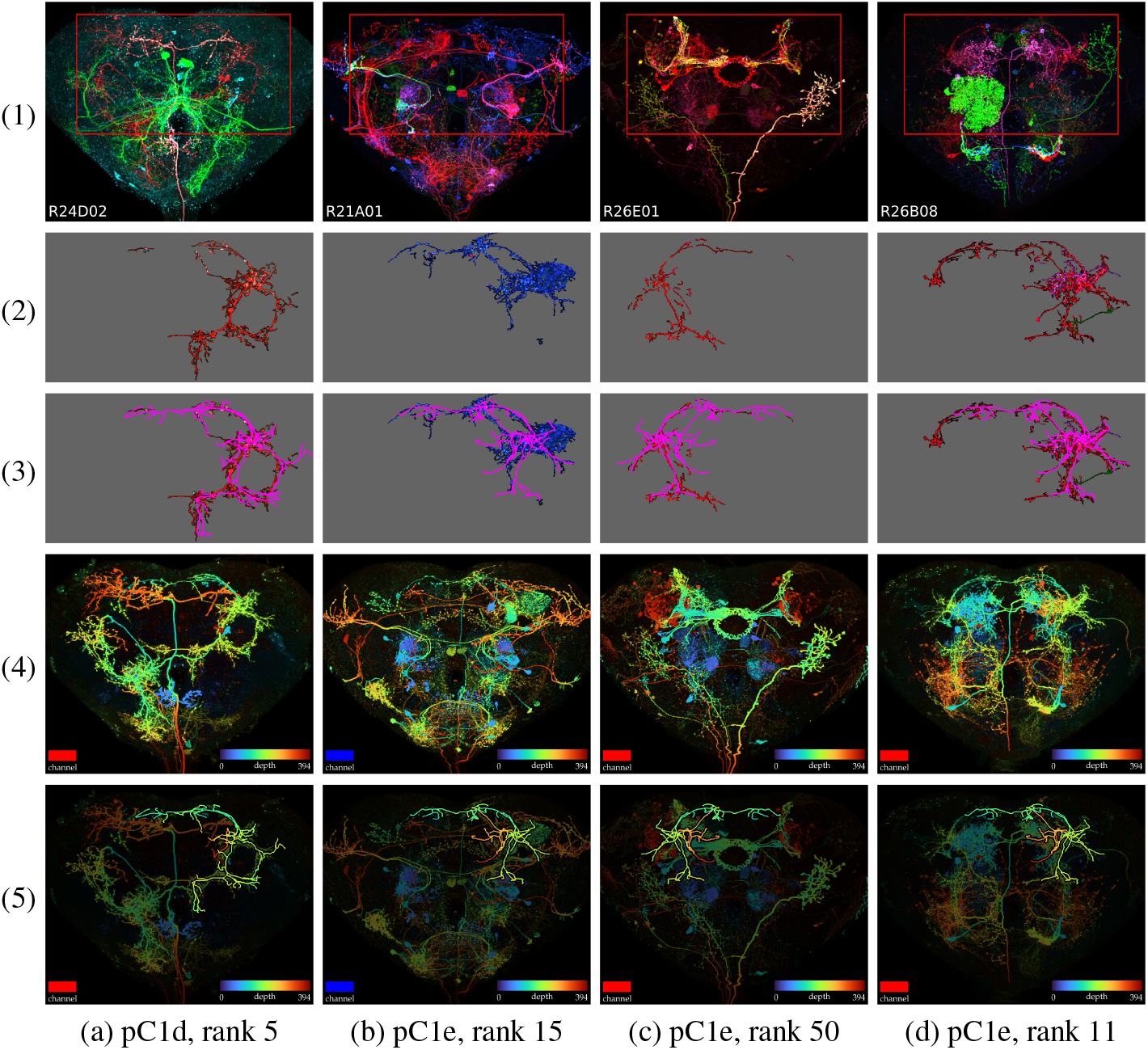
(a-c) Confirmed true positive and (d) convincing-looking albeit not confirmed PatchPerPixMatch hits for the target neurons pC1d and pC1e. The rows display the following from top to bottom: (1) Maximum intensity projection (MIP) of MCFO acquisition. (2) Section within the red rectangle from (1) masked by selected PatchPerPix fragments, (3) overlaid with skeletonized target, (4) Color-depth MIP [11] of best-matching channel, and (5) the same dimmed and overlaid with color-depth projection of the target. In (a), the target neuron can easily be spotted in rows 1, 4 and 5, whereas in (b) and (c), the PatchPerPixMatch fragment selection is required to reveal respective hits, as shown in rows 2 and 3.

By visual inspection, we found that many of the MCFO acquisitions that rank higher than those of known matching lines appear to contain visually plausible matches (cf. Figure 5d). However, their respective lines have not been subjected to split line creation and hence cannot be confirmed to be true matches. On the other hand, some high-ranking MCFO acquisitions appear to be false hits upon visual inspection, some of which can be attributed to false mergers of densely clustered neurons in the Patch-PerPix segmentation (cf. Figure 6a). Another potential cause for false hits are neurons with similar morphology, large fractions of which may get covered by high-scoring fragments. Figures 6b and 6c show respective false hits for an extreme case of similar morphologies, namely of the neurons pC1d and pC1e.

**Fig. 6:**
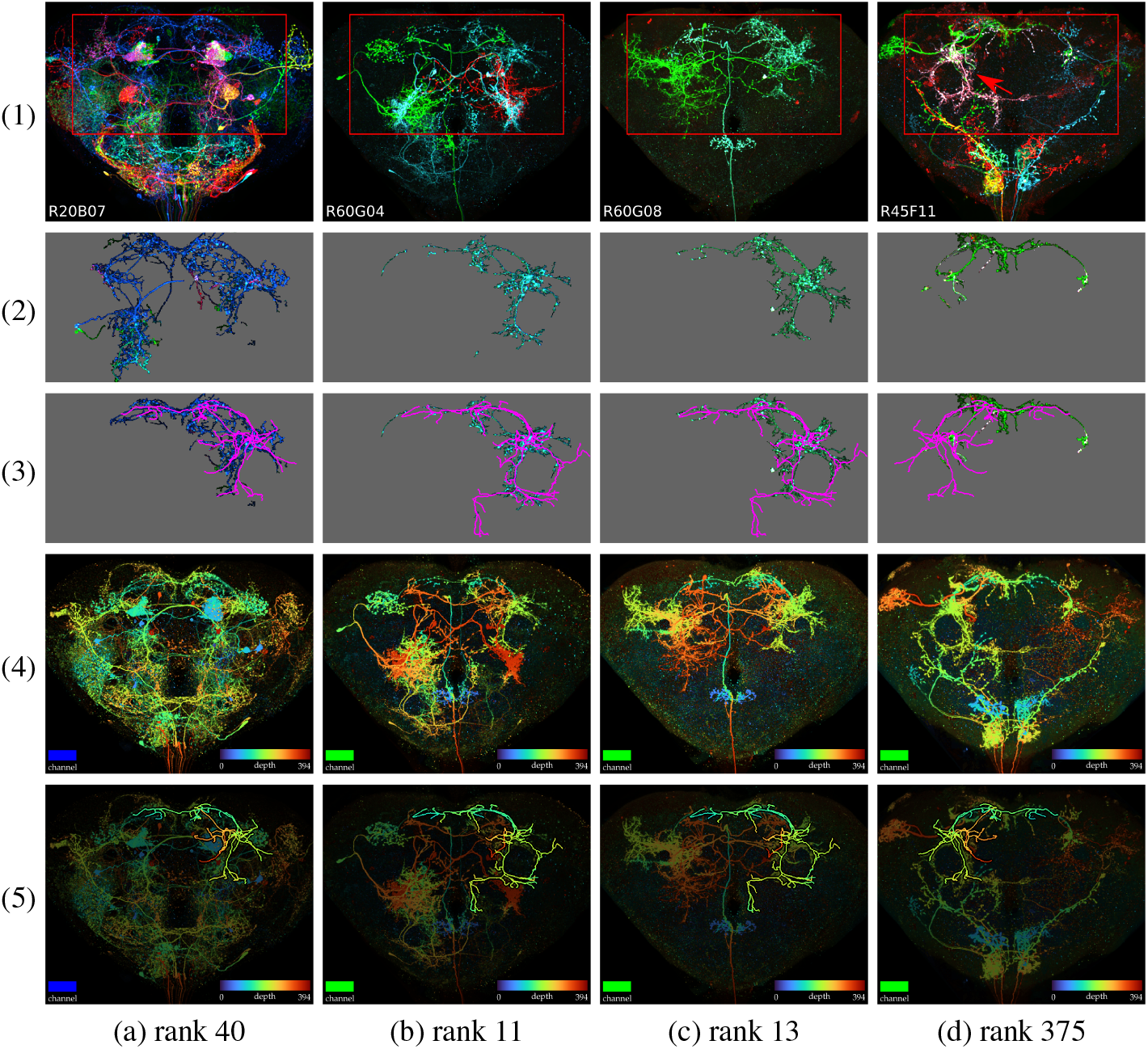
Failure cases / sources of error for PatchPerPixMatch. (a) False hit for the target neuron pC1e caused by false merge of densely clustered neurons. (b*c) False hits for the target neuron pC1d in lines that are known to exclusively contain pC1e: While most of the target neuron pC1d is covered by high-scoring fragments (resulting in high PatchPerPixMatch scores and ranks), the distinctive arbor of pC1d as compared to pC1e (cf. Figure 4a) is left uncovered. (d) False miss of the target neuron pC1e in an MCFO acquisition where pC1e overlaps with another neuron of different color, causing an observed shift in color. (For a row-by-row description of the visualization, see Figure 5.)

Some known matches missed by PatchPerPixMatch can be attributed to drastic shifts in color we empirically observe within some neurons in MCFO acquisitions (cf. Figure 6d). Such shifts may occur in case of overlap with a second neuron with (partly) similar morphology yet different color. Future work will investigate if a modified color difference penalty better handles these cases.

False misses may in general also be caused by the stochastic nature of MCFO labelling, due to which sets of MCFO acquisitions for a GAL4 line are not guaranteed to cover the complete respective GAL4 expression pattern [10]. Hence a neuron labelled in the GAL4 expression pattern may be missing in the respective MCFO image data as opposed to being missed by the PatchPerPixMatch search.

### Run-time Analysis and Code Availability

The average run-time for a PatchPerPix-Match search for an individual target morphology over all ~20,000 segmented MCFO acquisitions was 116 min on a single 3.0 GHz core.

Our code is available at *https://github.com/Kainmueller-Lab/PatchPerPixMatch*.

## 4 Visualization of PatchPerPixMatch Search Results

Our qualitative analysis reveals, apart from typical sources of error, which kinds of visualization of PatchPerPixMatch hits provide helpful information for the end user to effectively filter out false hits and increase the success rate of split-GAL4 line creation. Figure 7 depicts our thus-derived visualization for an exemplary match of a hemibrain neuron morphology in a GAL4 MCFO acquisition. It shows a PatchPerPixMatch result for a neuron that is easily visible in all panels of the visualization, for the purpose of clearly conveying our visualization.

**Fig. 7:**
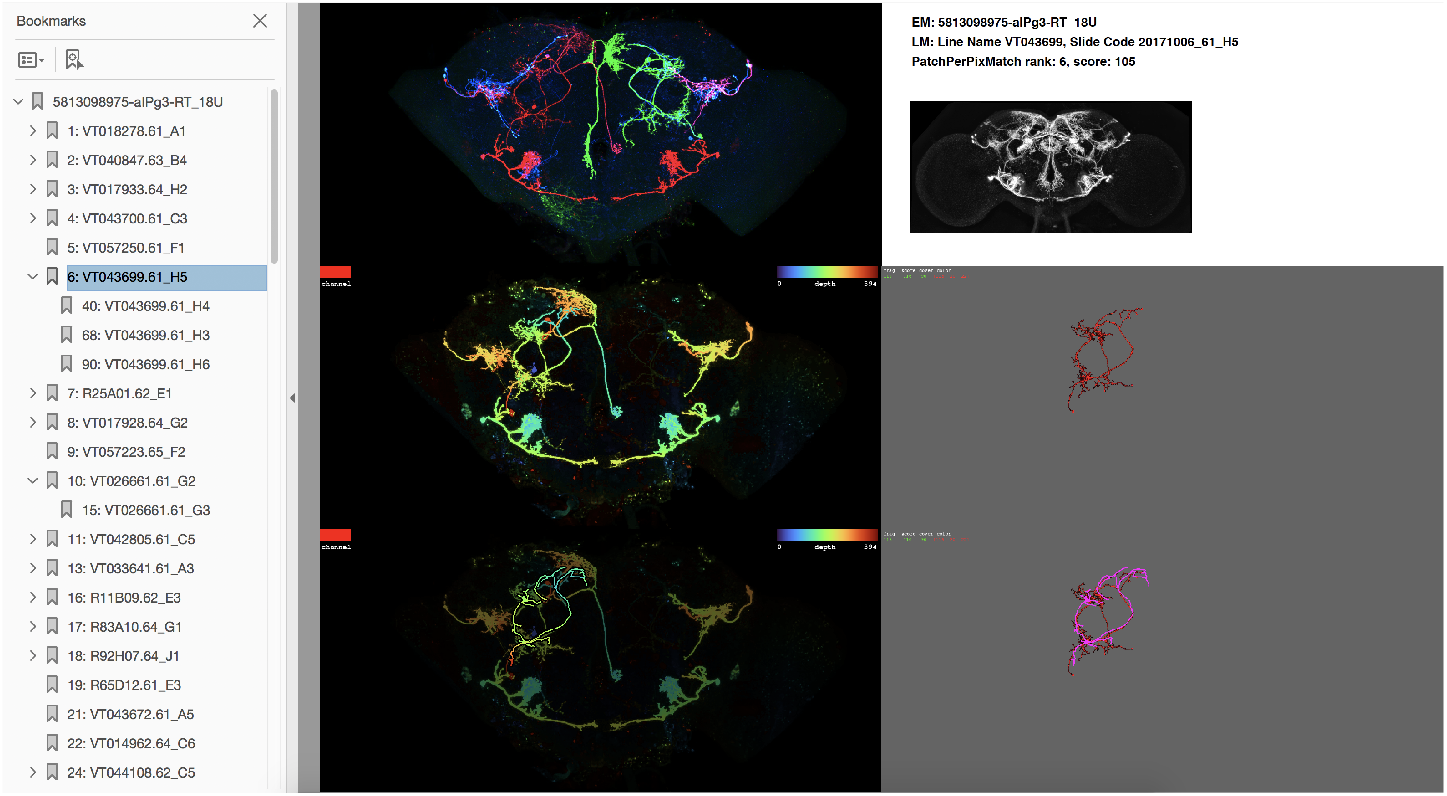
Our proposed visualization of a PatchPerPixMatch search result is composed of: **Top left image:** Maximum intensity projection (MIP) of the MCFO sample (all three neuron signal channels). **Mid left image:** Color-depth MIP (i.e. projection image with depth encoded as color) of best-matching channel. The channel color is indicated in the top left corner of the image. Color depth encoding is specified in the top right corner of the image. **Bottom left image:** Color-depth MIP of best-matching channel, dimmed and overlaid with color-depth projection of target neuron morphology. **Top right panel:** hemibrain ID of the target neuron, line- and sample ID of the MCFO sample, PatchPerPixMatch rank (among all ~20,000 searched MCFO acquisitions) and PatchPerPixMatch score, as well as a MIP of the full expression pattern of the GAL4 line. The latter is supposed to give a sense of the respective density. The PatchPerPix-Match score is defined as 100 *E*(**x**) to yield a positive score (higher is better) with negligible decimal places. **Mid right image:** MCFO MIP masked by best-matching PatchPerPix fragments. **Bottom right image:** MCFO MIP masked by best-matching PatchPerPix fragments (pruned if the match stems from a pruned run), overlaid with hemibrain body in magenta. **Left side-panel:** pdf bookmarks: Matches for a target morphology are sorted by PatchPerPixMatch ranks, and visualizations of the best 150 matches are assembled into a pdf document, with bookmarks that allow quick access via short summaries of matches (rank: line. sample). If top-150 matches occur in multiple samples of the same line, they are sorted in directly behind the best match of a line. This Figure is best viewed with zoom / on large screen.

If the sought target neuron is clearly visible in the maximum intensity projection (MIP) of the MCFO acquisition, then this is considered the best case scenario where split-GAL4 line creation can be done with high confidence. If an MCFO sample labels a higher number of neurons and the target thus cannot be spotted easily by eye in the respective MCFO MIP, it might still be visible in the color-depth MIP panels of the best-matching MCFO channel (as also employed in [11]), which exhibit reduced neuron density and furthermore enable direct comparison of 3d location between target and MCFO fragments. However, if the target is barely visible even in the channel color-depth MIP panels due to faintness or substantial occlusion by other neurons in the same channel, the panels showing MIPs of the fragment-masked MCFO acquisition (with and without the overlaid target) may still reveal a clean hit. These panels may also show more clearly than all others if the potential match misses to cover a characteristic part of the target, as seen e.g. for the false hits of the pC1d neuron in Figure 6b and 6c.

We created 30,000 pdf documents, one for each target neuron, visualizing the top 150 PatchPerPixMatch ranks as described in Figure 7. These pdfs are available for download at *https://pppm.janelia.org*. To facilitate annotation efforts, we also provide a spreadsheet for each target neuron morphology, where each row contains the GAL4 line name, MCFO slide code, and PatchPerPixMatch rank (among all ~20,000 searched MCFO acquisitions) as contained in the respective PDF, sorted by rank.

## 5 Discussion and Conclusion

We have developed PatchPerPixMatch, a fully automated method for efficient 3d search of Drosophila neuron morphologies in light microscopy imagery. PatchPerPixMatch is based on automatic neuron reconstructions, and features built-in robustness to respective segmentation inaccuracies. We release search results for over ~30,000 neuron morphologies in more than ~20,000 light microscopy acquisitions of Gen1 GAL4 line MCFO Drosophila brains. This resource serves to complement the 2d projection-based search results from [11,1].

PatchPerPixMatch is applicable to target neuron morphologies from any source, as long as the target is registered to the same reference coordinate frame as the segmentation fragments. Note, if a target neuron morphology stems from light microscopy as opposed to EM, like e.g. the neuron morphologies available in the *FlyCircuit* database [3], the step of pruning small branches of the target neuron (cf. *Target Point Influence* described in Sec. 2.1) may be dispensable. Concerning the suitability of PatchPerPix-Match for user-conducted searches for further target neuron morphologies in Gen1 GAL4 line MCFO brain acquisitions from [10], we expect near-interactive run-times (<10 minutes) on off-the-shelf workstations (3 GHz, 16 cores), deduced from an average single-threaded run-time of 116 minutes as measured in our study.

PatchPerPixMatch includes components that are, to various degrees, learnt from data: PatchPerPix [9] is based on a deep neural network trained for brain neuron instance segmentation, and NBLAST [6] similarity scoring is based on a histogram of atomic features observed for sets of neuron morphologies that include known matches. Consequently, generalizability to new data sources, which potentially follow distributions different from the ones employed in this work, needs to be assessed in respective further studies. E.g., while PatchPerPixMatch is technically not limited to the Drosophila brain, but can also be applied to the ventral nerve cord as is, respective potential distribution differences and their effect on PatchPerPixMatch performance remain to be assessed in future work.

Our released results are based on segmentation fragments obtained with PatchPerPix [9]. Technically, PatchPerPixMatch can take any automated segmentation result of the image data to be searched as input. However, respective segmentation accuracy, and, more specifically, the ratio of false merge- vs. false split segmentation errors, may impact PatchPerPixMatch performance. Future work will aim at end-to-end learning of neuron segmentation plus matching, where we will leverage a set of known matches on top of manual neuron segmentations as training data. Beyond replacing empirically determined parameter values in PatchPerPixMatch with machine-learnt values, such end-to-end learning may help gauge automated segmentation with PatchPerPix [9] towards an optimal balance of split- and merge errors.

Further future work will provide a quantitative comparison of PatchPerPixMatch with the heuristic, 2d projection-based method of Otsuna et al. [11,1], as well as with directly using NBLAST scores [6] to rank PatchPerPix fragments, in terms of their efficiency and effectiveness for finding GAL4 lines known to contain a target morphology. Last but not least, pre-computed PatchPerPixMatch search results will be integrated into the browser-based neuron search tool *NeuronBridge* [1] in a future release.

## Acknowledgments

We wish to thank Ruchi Parekh and the Janelia Connectome Annotation Team for providing pre-publication information on known matching GAL4 lines for a series of target neurons reconstructed from the hemibrain. We wish to thank Hideo Otsuna for providing a select list of ~30,000 hemibrain body IDs as well as their skeletons.

P.H., L.M. and D.K. were funded by the Max Delbrueck Center for Molecular Medicine, and supported by the HHMI Janelia Visiting Scientist Program. P.H. was funded by the MDC-NYU exchange program and HFSP grant RGP0021/2018-102.

## A Resources

**Fig. 8:**
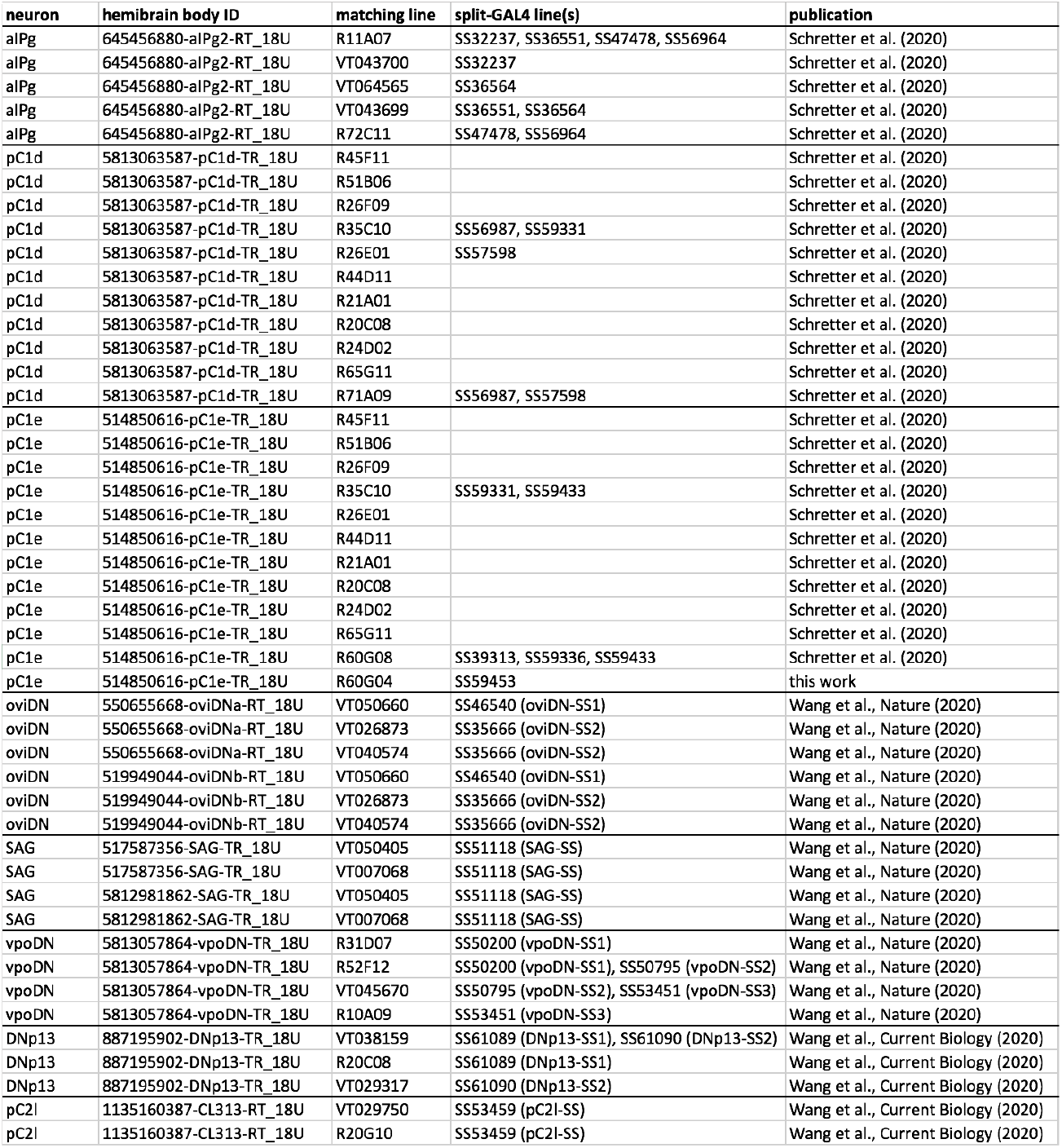
Known matching lines used in this work, and their origin.

## B Quantitative Evaluation on pC1d and pC1e Neurons

**Fig. 9:**
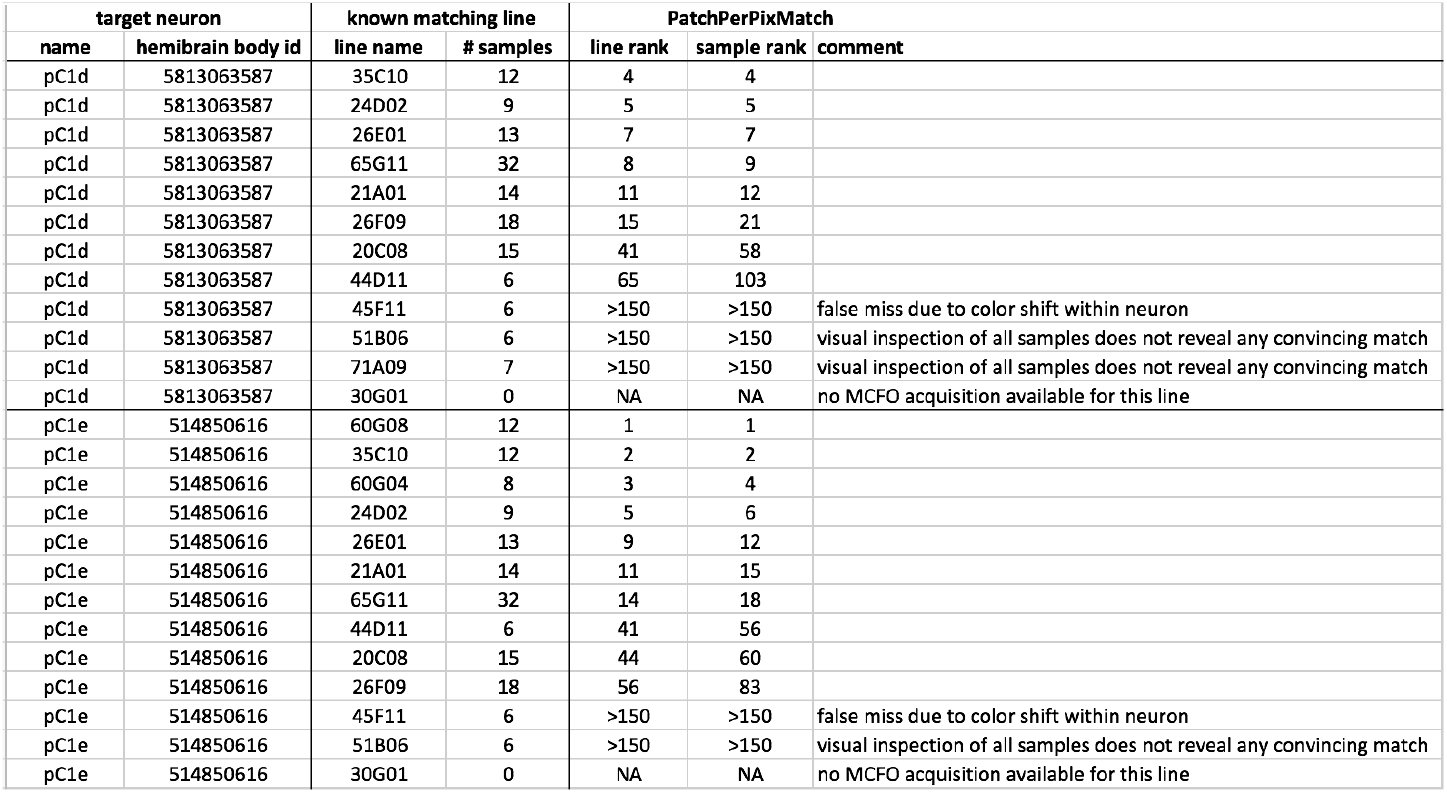
PatchPerPixMatch line- and sample ranks for known matching lines for the pC1d and pC1e target neurons.

**Fig. 10:**
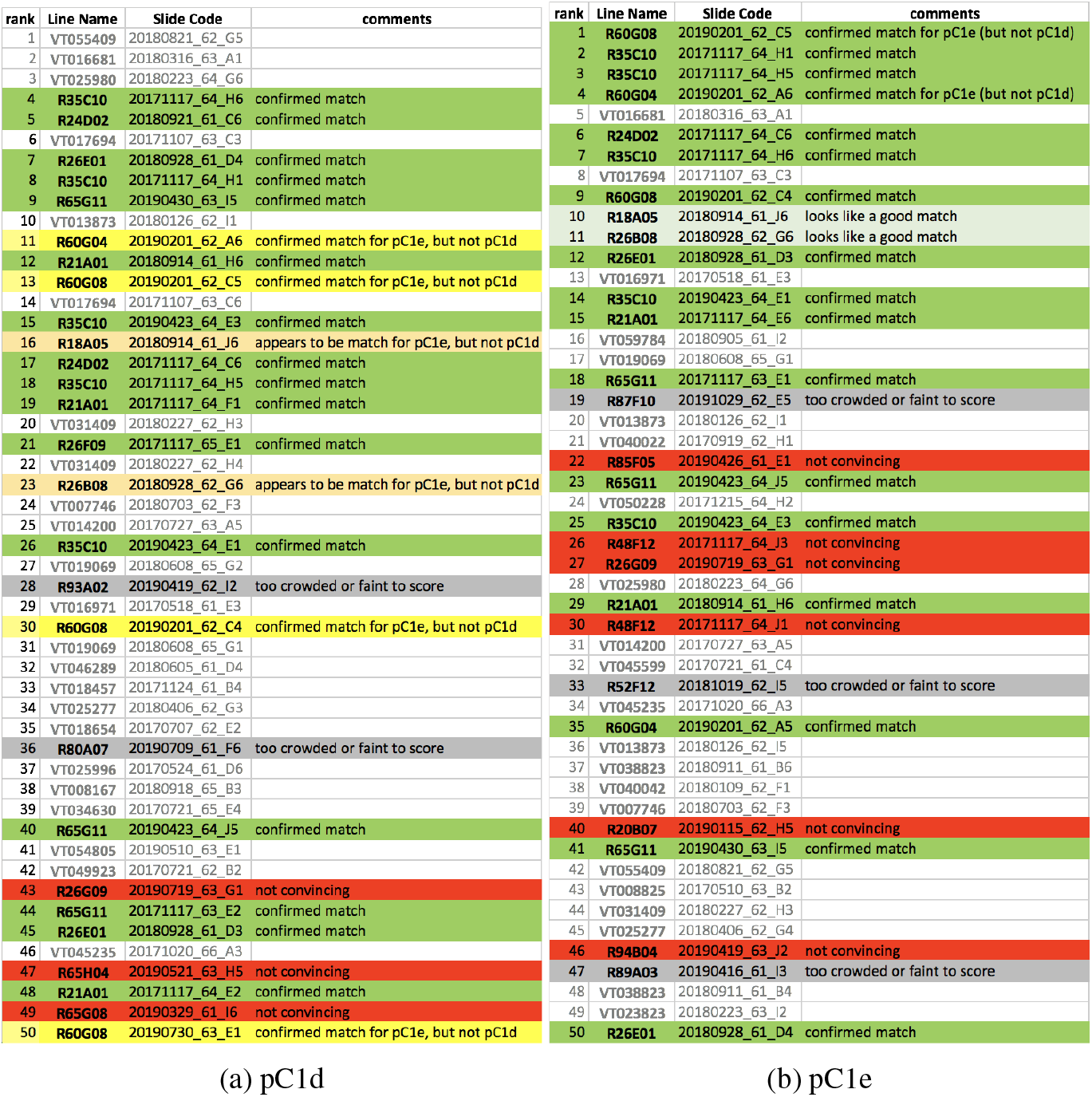
Annotated first 50 PatchPerPixMatch hits for the pC1d and pC1e target neurons. Confirmed matches for pC1d stem from an exhaustive behavioral screen over all R lines [17]. To this effect the set of known matching R lines for pC1d exhibits a level of completeness. Completeness of known matching lines allows for determining false PatchPerPixMatch hits. To this end we ignored the VT line hits here.

